# Impact of reinfection with SARS-CoV-2 Omicron variants in previously infected hamsters

**DOI:** 10.1101/2022.08.30.505966

**Authors:** Nozomi Shiwa-Sudo, Yusuke Sakai, Naoko Iwata-Yoshikawa, Shinji Watanabe, Souichi Yamada, Yudai Kuroda, Tsukasa Yamamoto, Masayuki Shirakura, Seiichiro Fujisaki, Kaya Miyazaki, Hideka Miura, Shiho Nagata, Shuetsu Fukushi, Ken Maeda, Hideki Hasegawa, Tadaki Suzuki, Noriyo Nagata

**Affiliations:** Department of Pathology, National Institute of Infectious Diseases, Tokyo, Japan; Research Center for Influenza and Respiratory Viruses, National Institute of Infectious Diseases, Tokyo, Japan; Department of Virology I, National Institute of Infectious Diseases, Tokyo, Japan; Department of Veterinary Science, National Institute of Infectious Diseases, Tokyo, Japan

## Abstract

The diversity of SARS-CoV-2 mutations raises the possibility of reinfection of individuals previously infected with earlier variants, and this risk is further increased by the emergence of the B.1.1.529 Omicron variant. In this study, we used an *in vivo*, hamster infection model to assess the potential for individuals previously infected with SARS-CoV-2 to be reinfected with Omicron variant and we also investigated the pathology associated with such infections. Initially, Syrian hamsters were inoculated with a lineage A, B.1.1.7, B.1.351, B.1.617.2 or a subvariant of Omicron, BA.1 strain and then reinfected with the BA.1 strain 5 weeks later. Subsequently, the impact of reinfection with Omicron subvariants (BA.1 and BA.2) in individuals previously infected with the BA.1 strain was examined. Although viral infection and replication were suppressed in both the upper and lower airways, following reinfection, virus-associated RNA was detected in the airways of most hamsters. Viral replication was more strongly suppressed in the lower respiratory tract than in the upper respiratory tract. Consistent amino acid substitutions were observed in the upper respiratory tract of infected hamsters after primary infection with variant BA.1, whereas diverse mutations appeared in hamsters reinfected with the same variant. Histopathology showed no acute pneumonia or disease enhancement in any of the reinfection groups and, in addition, the expression of inflammatory cytokines and chemokines in the airways of reinfected animals was only mildly elevated. These findings are important for understanding the risk of reinfection with new variants of SARS-CoV-2.

**IMPORTANCE:** The emergence of SARS-CoV-2 variants and the widespread use of COVID-19 vaccines has resulted in individual differences in immune status against SARS-CoV-2. A decay in immunity over time and the emergence of variants that partially evade the immune response can also lead to reinfection. In this study, we demonstrated that, in hamsters, immunity acquired following primary infection with previous SARS-CoV-2 variants was effective in preventing the onset of pneumonia after reinfection with the Omicron variant. However, viral infection and multiplication in the upper respiratory tract were still observed after reinfection. We also showed that more diverse nonsynonymous mutations appeared in the upper respiratory tract of reinfected hamsters that had acquired immunity from primary infection. This hamster model reveals the within-host evolution of SARS-CoV-2 and its pathology after reinfection, and provides important information for countermeasures against diversifying SARS-CoV-2 variants.

## INTRODUCTION

After the emergence of the severe acute respiratory syndrome coronavirus 2 (SARS-CoV-2), at the end of 2019, various Variants of Concern (VOC) emerged, including the B.1.1.7 (Alpha), B.1.351 (Beta) and B.1.617.2 (Delta) strains, allowing COVID-19 infection to spread and persist worldwide (1). At the end of November 2021, the World Health Organization designated a variant of SARS-CoV-2, B.1.1.529, as a VOC (2). The B.1.1.529, Omicron variant, has 30 amino acid substitutions, three deletions, and one insertion site in the spike region compared with the ancestral, Wuhan, SARS-CoV-2 strain, including 15 mutations in the receptor binding region (3). The emergence of Omicron strains with high transmission capacity has changed the infection risk situation for the current COVID-19 pandemic (3, 4). Although several studies indicate that Omicron variants cause less severe disease than the Delta variant (5), it is clear that an individual infected with an earlier VOC or Omicron strain is still at risk of reinfection with the next variant, including Omicron subvariant (6). In addition, infection with Omicron is considered primarily as an upper respiratory tract infection. This differs from the pathophysiology associated with the earlier VOC, which tended to cause lower respiratory tract infection (7–10). Thus, there is increasing concern about the efficacy of immunity of previously infected individuals against the new variants and impact of immunopathology due to reinfection of SARS-CoV-2 (11, 12).

The Syrian hamster is more susceptible to SARS-CoV-2 than other animal species (13). Weight loss, clinical signs, pathology, and immune response can be used as indicators of viral infection in hamsters, making them a useful small animal model for development of vaccines and antiviral agents for COVID-19 (13–15). Several research groups have also used this animal model to investigate the phenotype of mutant viruses by testing for changes in infectivity, infectiousness, and antigenicity (16, 17). Hansen et al used a hamster model to conduct reinfection experiments with a homologous, ancestor strain, WA1, and heterologous B.1.1.7 (Alpha) and B.1.351 (Beta) SARS-CoV-2 variants to determine the transmission through reinfection of asymptomatic individuals (18). Reinfection leads to SARS-CoV-2 replication in the upper respiratory tract with the potential for virus shedding, suggesting the risk of transmission through reinfected asymptomatic individuals. On the other hand, another group showed that prior infection with the WA1 strain prevented Delta variant transmission to naïve hamsters (19). In this study, therefore, we used a hamster model to i) evaluate the risk of reinfection with Omicron BA.1 strain, following infection with prior VOC strains, and ii) determine the risk of reinfection with Omicron subvariants in individuals first infected with the Omicron BA.1 strain.

## RESULTS

### Primary infection of hamsters with SARS-CoV-2 variants

Five isolates of SARS-CoV-2 including the ancestor strain from lineage A, and four isolates of the VOC (B.1.1.7, B.1.351, B.1.617.2 and B.1.1.529; BA.1.18 [referred to as BA.1]; Table 1) were used for primary inoculation of Syrian hamsters as depicted in Figure 1A. Because hamsters are highly susceptible to SARS-CoV-2, primary infection was conducted by intranasal inoculation of a low dose of virus (1.0×10^3^ TCID_50_ in 8 μL of Dulbecco’s Modified Eagle Medium (DMEM)), which reached the local upper respiratory tract, and induced seroconversion. Five weeks after primary infection, the lower respiratory tract of animals was then reinfected with a higher dose (1.0×10^4^ TCID_50_ in 50 μL) of virus.

**TABLE 1.** SARS-CoV-2 variants in this study

| Pango | Sub | Strain | GISAIID Accession | Note |
| --- | --- | --- | --- | --- |
| lineage | lineage | (Simplified name) | no. of original |  |
| (WHO |  |  | isolate |  |
| label) |  |  |  |  |
| A |  | hCoV-19/Japan /TY-WK-521/2020 (WK-521) | EPI_ISL_408667 |  |
| B.1.1.7 |  | hCoV-19/Japan/QK002/2020 | EPI_ISL_768526 |  |
| (Alpha) |  | (QK002) |  |  |
| B.1.351 |  | hCoV-19/Japan/TY8-612- | EPI_ISL_1123289 |  |
| (Beta) |  | P1/2021 (TY8-612) |  |  |
| B.1.617.2 |  | hCoV-19/Japan/TY11-927- | EPI_ISL_2158617 |  |
| (Delta) |  | P1/2021 (TY11-927) |  |  |
| B.1.1.529 | BA.1.18 | hCoV-19/Japan/TY38- | EPI_ISL_7418017 | BA.1 + |
| (Omicron) |  | 873P0/2021 (TY38-873) |  | ORF1a:T1822I |
| B.1.1.529 | BA.1.1 | hCoV-19/Japan/TY38- | EPI_ISL_7571618 |  |
| (Omicron) |  | 871P0/2021 (TY38-871) |  |  |
| B.1.1.529 | BA.2 | hCoV-19/Japan/TY40-385- | EPI_ISL_9595859 |  |
| (Omicron) |  | P1/2022 (TY40-385) |  |  |
| B.1.1.529 | BA.2.3 | hCoV-19/Japan/TY40-816- | EPI_ISL_9595861 | BA.2 + |
| (Omicron) |  | P1/2022 (TY40-816) |  | S:A688V |

**FIG 1.**
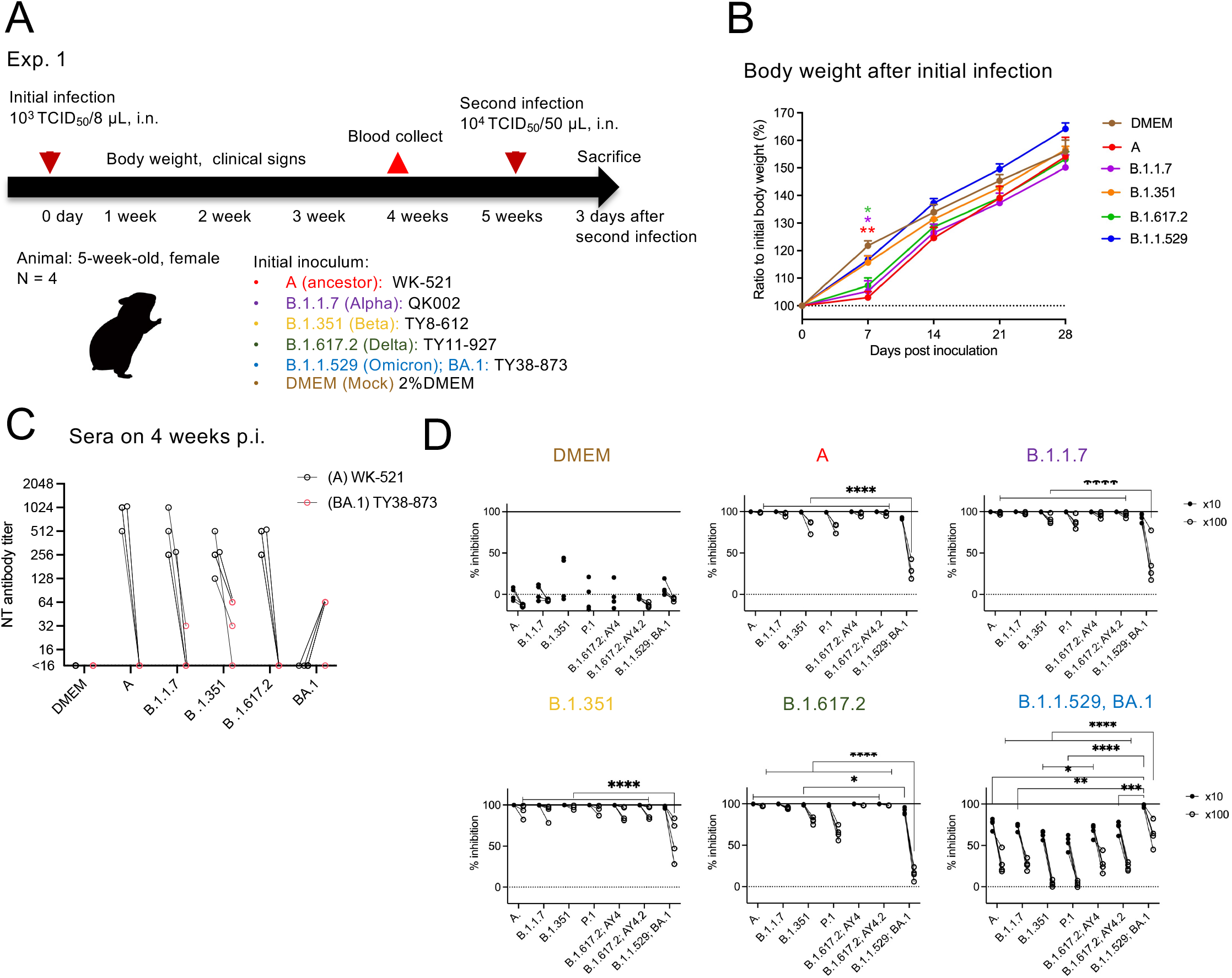
Experimental infection of hamsters with SARS-CoV-2 variants. Study design outlining experimental infections using 5-week-old female Syrian hamsters and SARS-CoV-2 **(A)**. Body weight curve for 4 weeks after primary infection with SARS-CoV-2 variants. Dunnett’s multiple comparison test compared with DMEM-inoculated animals (n = 3 or 4). *, P < 0.05; **, P < 0.01 **(B)**. Neutralizing (NT) antibody titers against WK-521 (lineage A, black circle) or TY38-873 (lineage B.1.1.529, BA.1, **(C,** red circle)) strains in the sera from hamsters at 4 weeks post-inoculation (p.i.). Data from the same animal are connected with lines. *, P < 0.05 by Dunn’s multiple comparison test following the Kruskal–Wallis test **(C)**. Blocking activities of hamster sera between ACE2 and SARS-CoV-2 spike was tested by the ACE2 binding inhibition ELISA using a V-Plex COVID-19 ACE2 neutralization kit (K15570U, Meso Scale Discovery) (n = 3 or 4). Each dot represents data from an individual animal. Sera were diluted 1:10 (black dot) or 1:100 (black circle) for the analysis. Data from the same animal are connected with lines. *, P < 0.05; **, P < 0.01; ***, P < 0.001; ****, P < 0.0001 by Sidak’s multiple comparison test following two-way ANOVA **(D)**.

After the primary inoculation, animals infected with isolates from lineage A, B.1.1.7, and B.1.617.2 showed significantly lower weight gain rate than the mock-infection group (DMEM in Figure 1B, n = 3–4) in the first week post-inoculation (p.i.) but showed no differences by four weeks p.i. (Figure 1B). At four weeks p.i., all animals, except those inoculated with the B.1.1.529; BA.1 strain, showed high antibody titers against the ancestor strain, lineage A and poor neutralizing activities against B.1.1.529; BA.1 strain (Figure 1C). The B.1.1.529; BA.1 strain-inoculated animals had neutralizing antibodies (1:64) against the homologous strain, but no neutralizing activities against the ancestor strain (detection limit was 1:16). Sera were also used in a multiple assay for antibodies that block the binding of human ACE2 to the spike proteins from variants of SARS-CoV-2 (Figure 1D). Lower inhibition activity against the spike from the B.1.1.529; BA.1 variant was seen in the sera from the A-, B.1.1.7-, and B.1.617.2-inoculated animals than in the sera from the B.1.351- and B.1.1.529 especially when used 1:100 diluted sera; BA.1-infected animals. Significantly, poor inhibition activity against the earlier VOC spike forms was observed in sera from the B.1.1.529; BA.1-infected animals.

### Reinfection of hamsters with an Omicron variant

Five weeks after the primary inoculations, the B.1.1.529; BA.1 strain was inoculated into all animals. No obvious respiratory illness was seen in any reinfected animals in the three days after the second inoculation. The homologous reinfected hamsters (BA.1-BA1 group in Figure 2A) showed transient body weight loss at 1 day post second infection unlike the mock infected- and primary infected-animals (DMEM group and DMEM-BA.1 group, respectively) by day 3 (Figure 2A). Three days after the second inoculation, all animals were euthanized, under overdose anesthesia, and blood, nasal wash fluid, and lung samples were obtained. The reinfected groups (A-BA.1 and B.1.1.7-BA.1 groups) showed significantly higher lung weight/body weight ratios at 3 days p.i. (Figure 2B). Infectious virus was detected in the respiratory samples of some of the primary infected hamsters (DMEM-BA.1 group, one of four in the nasal wash fluid; three of four in the lung homogenate), but no infectious virus was recovered from any of the reinfected animals (Figure 2C; detection limit was 10^1.5^ TCID_50_/mL). Interestingly, high viral RNA levels were detected in the nasal wash fluid from almost all animals, even though infectious virus was not isolated from these animals (Figure 2D upper panels). By contrast, no virus-associated RNA was detected in the lungs of animals with prior homologous BA.1 infection (BA.1-BA.1 group), and much lower virus copy numbers of virus-related RNA were detected in the lungs of animals with prior heterologous SARS-CoV-2 infection compared with those in the lungs of animals of DMEM-BA.1 group (Figure 2D lower panels). Thus, both prior homologous and heterologous SARS-CoV-2 infection elicited lower immune protection in the upper respiratory tract than that in the lower respiratory tract in this animal model.

**FIG 2.**
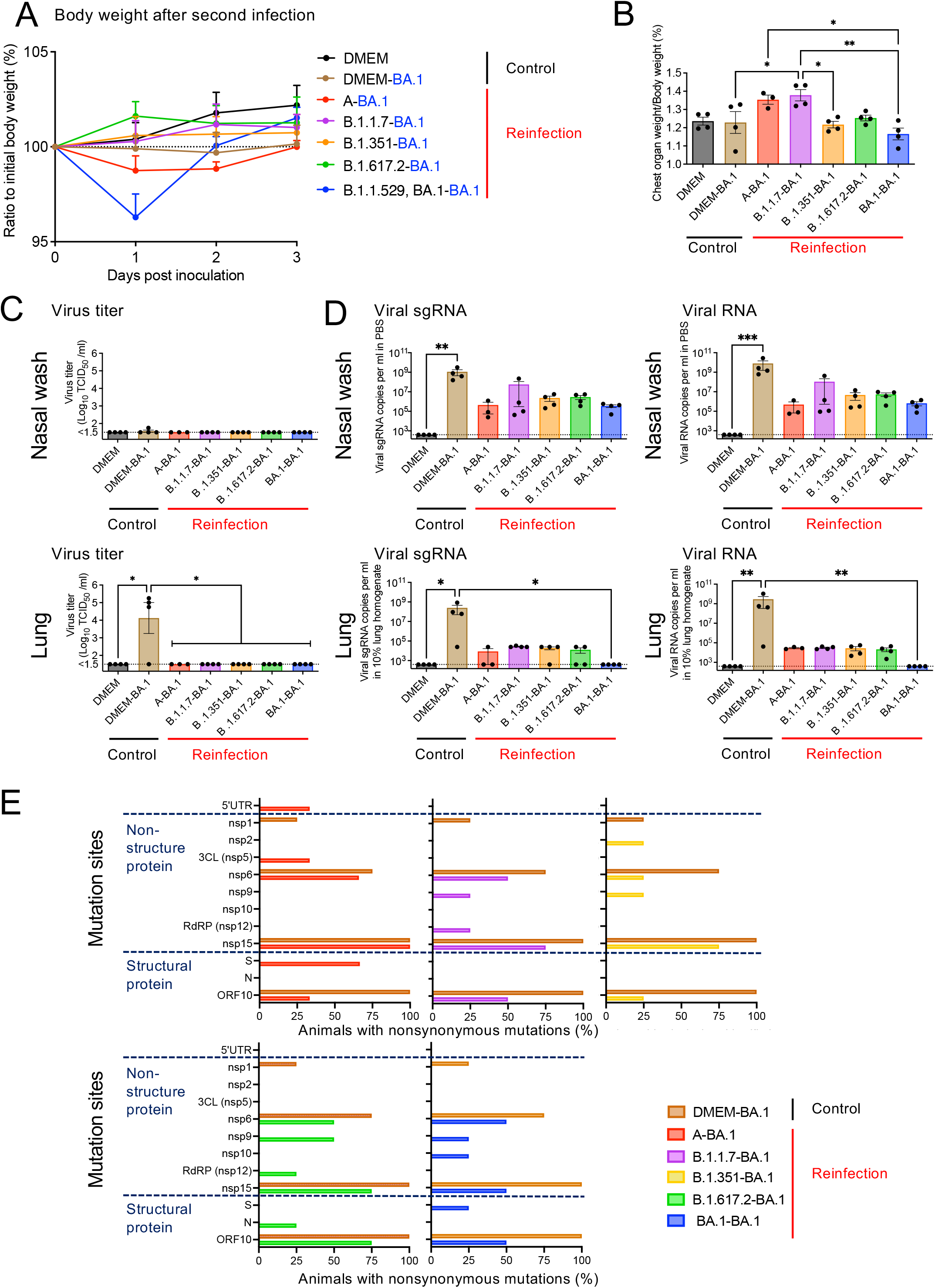
Reinfection of hamsters with an Omicron variant. Body weight curve during 3 days after second infection with an isolate of the TY38-873, B. 1.1.529 (Omicron) subvariant BA.1 (n = 3 or 4) **(A)**. Bar graph showing the ratio of the chest organ weights including lungs, trachea, heart, and thymus per body weight at 3 days p.i. (n = 3 or 4) **(B)**. Bar graph showing virus titers **(C)** and virus-related RNA copies **(D)** in the nasal wash fluid (upper panels) and supernatant from 10% lung tissue homogenates (lower panels) (n = 3 or 4). *, P < 0.05; **, P < 0.01; ***, P < 0.001; by Dunn’s multiple comparison test following the Kruskal–Wallis test. Percentage of animals with mutations involving amino acid substitutions in each region **(E)**. The full-genome sequences of the Omicron BA.1 strains recovered from the nasal wash fluid samples (n = 3 or 4). Brown bars in each panel indicate the same data for the DMEM-BA.1 group.

The full-genome sequence of each of the Omicron BA.1 variants recovered from the nasal wash fluid samples was determined (Supplementary Table 1). Nsp15 (ORF1b: K2340T) and ORF10 (ORF10: V30L) nonsynonymous mutations appeared in all four hamsters whose primary infection was with BA.1. (Figure 2E, left). Three of the four animals also harbored variants with one or two amino acid substitutions in ORF1a region (nsp1:G180E and/or nsp6:L260F). The same within-host variants could be observed in the animals from the DMEM-BA.1 group. However, in the animals with prior homologous BA.1 infection (BA.1-BA.1 group), nonsynonymous mutations were unevenly distributed among individuals with low or high frequency mutations in nonstructural genes (nsp6:L37F, nsp6:L260F, nsp9:T67A, nsp10:Q36R, nsp15:K289T) and structural genes (S: M1del and ORF10: V30L). Mutations with greater diversity were also detected from animals subjected to prior heterologous SARS-CoV-2 infection (Supplementary Table 1, Figure 2E, right). Common nonsynonymous mutations were observed in the upper respiratory tract of BA.1-primary infected hamsters, but diverse mutations appeared in that of BA.1-reinfected hamsters.

### Pathology in the respiratory tract after reinfection

Histopathological changes in reinfected animals were determined. Three days after reinfection, pathological lesions consisted of mild to moderate rhinitis and focal broncho-interstitial pneumonia in the respiratory tract of the primary infection group (DMEM-BA.1 group, Figure 3A, second row). SARS-CoV-2 N antigen-positive cells were observed both in the respiratory and olfactory epithelium of the nasal cavity and in bronchiolar epithelium and alveolar epithelia of the lungs (Figure 3B).

**FIG 3.**
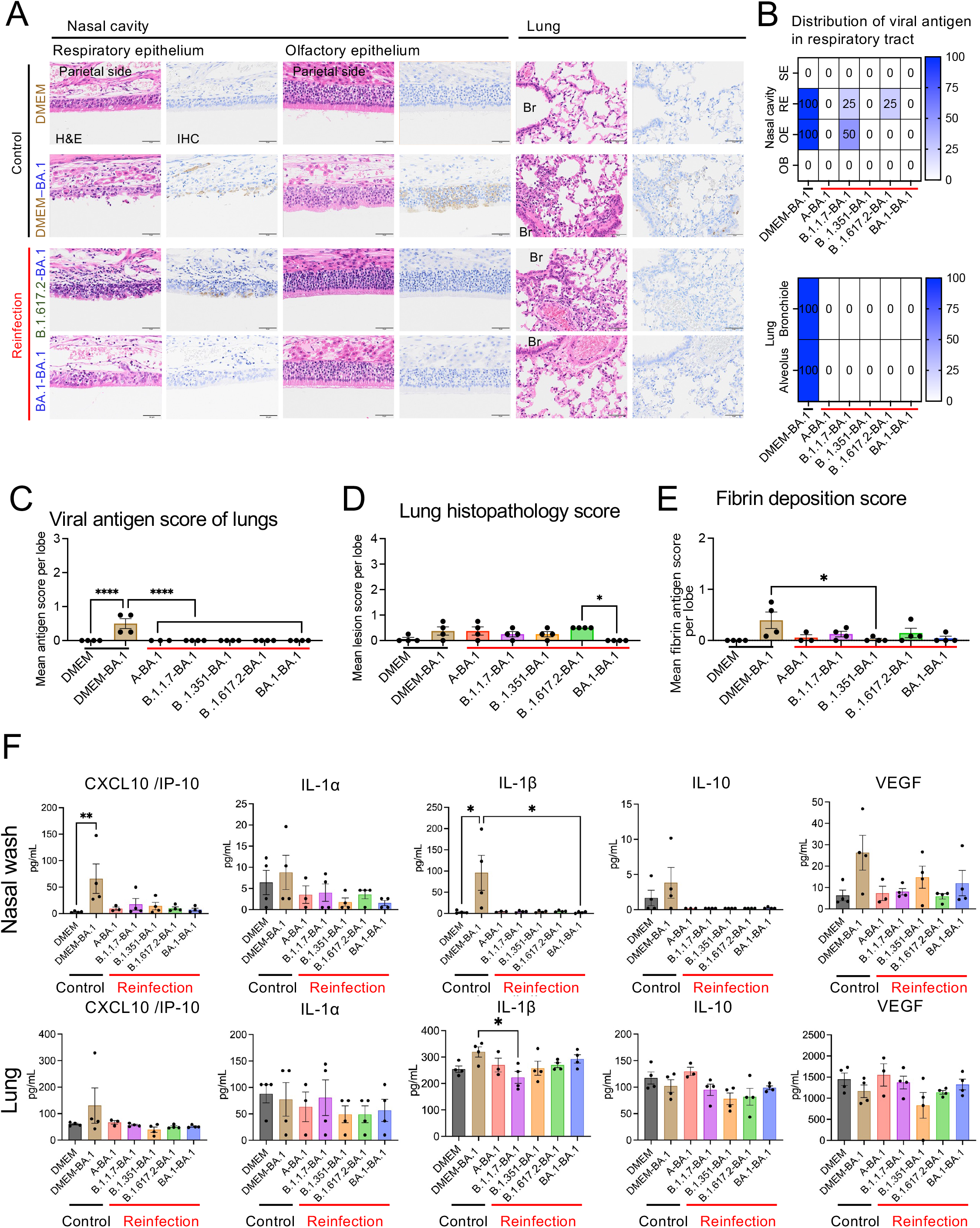
Histopathology and immunopathology of respiratory tract after reinfection with an Omicron BA.1 variant. Representative histopathological findings for the nasal cavity epithelia and lungs of hamsters 3 days after second infection with an isolate of TY38-873, B. 1.1.529 (Omicron) subvariant BA.1 (n = 4). H&E, hematoxylin and eosin staining; IHC, immunohistochemistry for SARS-CoV-2 antigen detection. Br, bronchi. Viral antigen-positive cells were mainly detected in the nasal epithelium on the cranial side. Scale bars, 50 μm **(A)**. Distribution of viral antigens in nasal cavity and lungs by immunohistochemistry. Heat map shows percentages of viral antigen-positive animals. SE; squamous epithelium, RE; respiratory epithelium, OE; Olfactory epithelium, OB; Olfactory bulb **(B)**. Viral antigen scores and pathological severity scores of lungs from hamsters (**C and D**). Fibrin deposition scores of lungs by immunohistochemistry. Four lung lobes were taken from each individual animal and scored to evaluate comprehensive pathological changes. *p < 0.05 by one-way ANOVA **(E)**. Cytokine and chemokine levels in the nasal wash fluid (upper panels) and supernatant from 10% lung tissue homogenates (lower panels) of hamsters at 3 days p.i. (n = 3 or 4). *, P < 0.05 by Dunn’s multiple comparison test following the Kruskal–Wallis test **(F)**.

In the homologous infection group (BA.1-BA.1), the lungs were hardly damaged by the primary infection and the onset of pneumonia, following reinfection, was prevented (Figure 3A-D). By contrast, the heterologous reinfection groups showed moderate lymphocyte infiltration in the respiratory area of nasal cavity in the absence of viral antigens, except in a few animals from the B.1.1.7-BA.1 group (two of four animals) and the B.1.617.2-BA.1 group (one of four animal); viral antigens were detected in the epithelia of these animals and marked lymphocytic infiltration was observed from the lamina propria into the epithelium with or without multilayering of respiratory epithelial cells (regeneration) in the nasal cavity (Supplementary Figure 1 and Figure 3A). The lungs from the heterologous reinfection groups showed small clusters of lymphocytes, plasma cells, and macrophages around the bronchioles and blood vessels in the absence of detectable viral antigens (Supplementary Figure 1, Figure 3A, B, and C). Bronchiolar regeneration was also observed in the reinfection groups. In particular, the epithelial regeneration was more pronounced in the lungs of the A-BA.1 group than in the lungs of other groups, indicating the severe damage caused by the primary infection with the lineage A strain (Supplementary Figure 1). In the B.1.617.2-BA.1 group, lymphocyte and macrophage infiltration was predominantly seen around the bronchioles, as reflected in the lung tissue score (Figure 3D). Fibrin deposition in the lungs is one of the main histopathological features of COVID-19 related acute pneumonia (20, 21). Fibrin deposition was often observed in lungs of the primary infected animals but in very few of the reinfected animals (Figure 3E).

High levels of cytokines and chemokines, including CXCL10/IP-10 and IL-1β, were observed in the nasal wash fluid of the primary infection group (DMEM-BA.1), but these were lower in reinfected animals (Figure 3F). Neither primary nor secondary infections with the BA.1 strain induced cytokine and chemokine profiles typically associated with pneumonia. Exacerbated cytokine production due to reinfection was not observed in either the upper or lower respiratory tract. Taken together, these data suggest that viral replication after reinfection was more strongly suppressed in the lower respiratory tract than in the upper respiratory tract.

### Experimental hamster infection with Omicron subvariants

Next, we evaluated the risk of reinfection with B.1.1.529, Omicron subvariants in hamsters first infected with the BA.1 strain. Hamsters were infected as above, using four isolates of the B.1.1.529 subvariants (BA.1, BA.1.1, BA.2 and BA.2.3; Table 1) were used for primary and/or secondary inoculation of Syrian hamsters as depicted in Figure 4A. After primary inoculation with the BA.1 strain, animals showed slightly lower weight gain rate than the control animals (n = 16) during the first week p.i. but no difference was evident by the second week p.i. (Figure 4B). At four weeks p.i., all BA.1 strain-inoculated animals showed seroconversion (Figure 4C). Clear antigenic differentiation between the BA.1 and BA.2 subvariants was shown in the sera from hamsters. While no neutralizing activity against the BA.2 strain was detected by a neutralization assay (Figure 4C), an assay measuring antibodies that block the binding of human ACE2 to spike proteins suggested the BA.1-infected hamster sera had inhibition activities against the spike from Omicron subvariants (Figure 4D).

**FIG 4.**
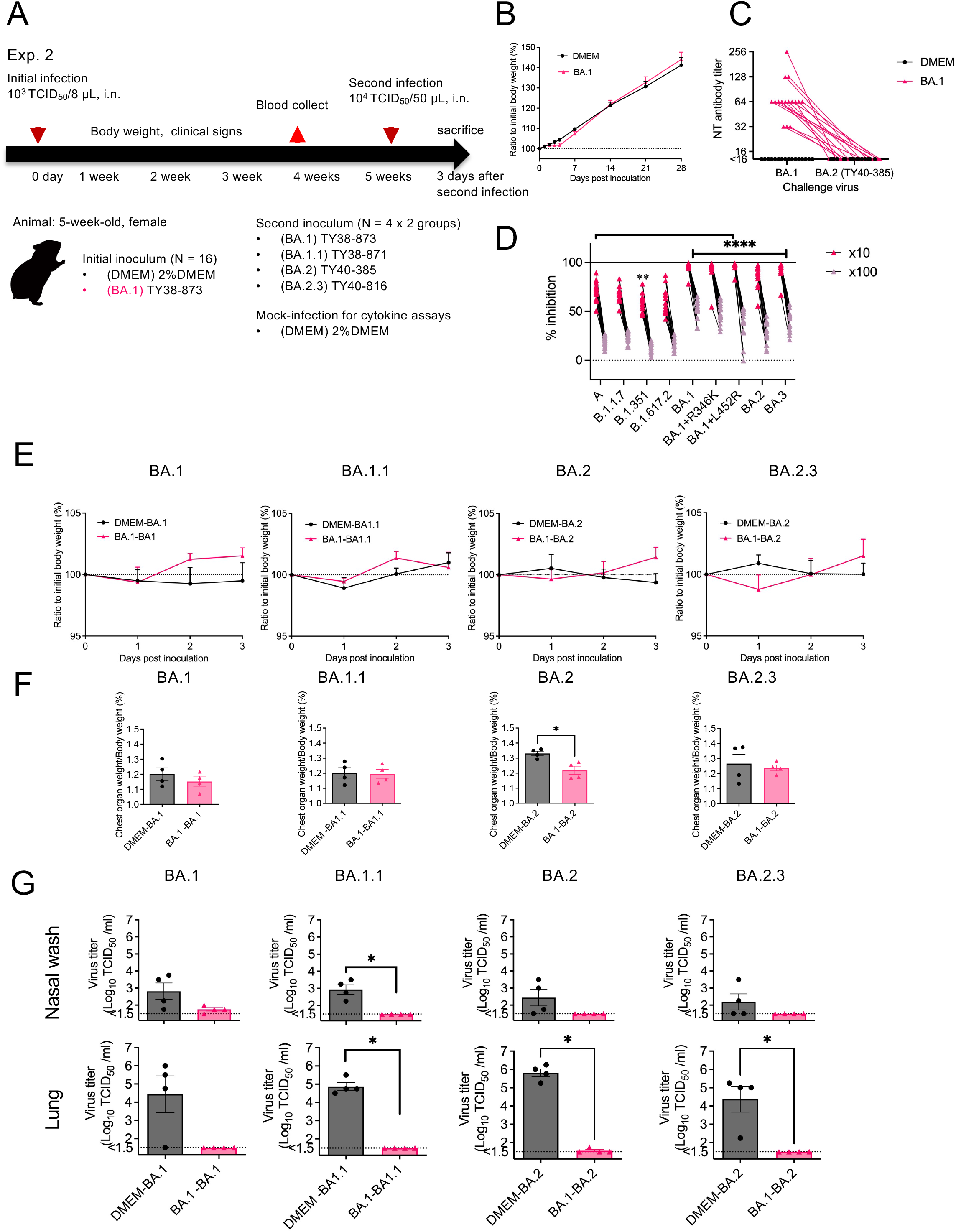
Experimental infection of hamsters with Omicron subvariants. Study design outlining experimental infections using 5-week-old female Syrian hamsters and Omicron subvariants **(A)**. Body weight curve for 4 weeks after primary inoculation with an isolate of TY38-873, B. 1.1.529 (Omicron) subvariant BA.1. Control groups were inoculated with cell culture medium (DMEM). No significant difference compared with DMEM-inoculated animals (n = 16) was detected by Sidak’s multiple comparison test following two-way ANOVA **(B)**. Neutralizing (NT) antibody titers against TY38-873 (BA.1, left dots and triangles) or TY40-385 (BA. 2, right dots and triangles) strains in the sera from hamsters on 4 weeks p.i. The dashed line indicates the limit of detection (<16). Each dot represents and triangles data from an individual animal (n = 16). Data from the same samples are connected with lines **(C)**. Blocking of interactions between ACE2 and SARS-CoV-2 spike was tested by ACE2 binding inhibition ELISA using a V-Plex COVID-19 ACE2 neutralization kit (K15586U, Meso Scale Discovery) (n = 16). Sera were diluted 1:10 or 1:100 for the analysis. **, P < 0.01; ****, P < 0.0001 by Sidak’s multiple comparison test following two-way ANOVA **(D)**. Body weight curve during 3 days after primary or reinfection with subvariants of B. 1.1.529. (n = 4). No significant differences were detected between the primary (black lines) and reinfection (red lines) groups by two-way ANOVA **(E)**. Bar graph showing the ratios of the weights of the chest organ including lungs, trachea, heart, and thymus to body weights at 3 days p.i *, P < 0.05 by Mann-Whitney test **(F)**. Bar graph of virus titers in the nasal wash fluid (upper panels) and supernatant from 10% lung tissue homogenates (lower panels) **(G)**. Dot line indicates detection limit. *, P < 0.05 by Mann-Whitney test., Black lines/bars indicate the primary infection groups and red lines/bars are from the reinfection groups (E-G).

Five weeks after the primary inoculation, B.1.1.529 subvariants (n = 4 per subvariant) were inoculated into the animals. No obvious respiratory illness was seen in any of the animals in the three days following the second inoculation. Body weight graphs did not show any significant difference between the primary and the reinfection groups (Figure 4E). Only animals infected with the BA.2 variant showed a significant difference in the lung/body weight ratio between primary and reinfection groups (Figure 4F). Infectious virus was detected from the respiratory samples of most hamsters after primary infection, but very low or no infectious virus was detected in samples from the reinfected animals (Figure 4G). However, infectious virus was detected in the nasal wash of three of the four animals in the BA.1-BA.1 reinfection group and the lungs of one of the four animals in the BA.1-BA.2 reinfection group. Despite the absence of infectious virus, high copy numbers of virus-associated RNA were detected in nasal lavage fluid from both homologous and heterologous reinfected animals. By contrast, significantly lower copy numbers of virus-associated RNA were detected in the lungs of reinfected animals than in those of the primary infected group (Figure 5A). Thus, animals with prior infection with BA.1 showed low protection of the upper respiratory tract against reinfection with the subvariants.

**FIG 5.**
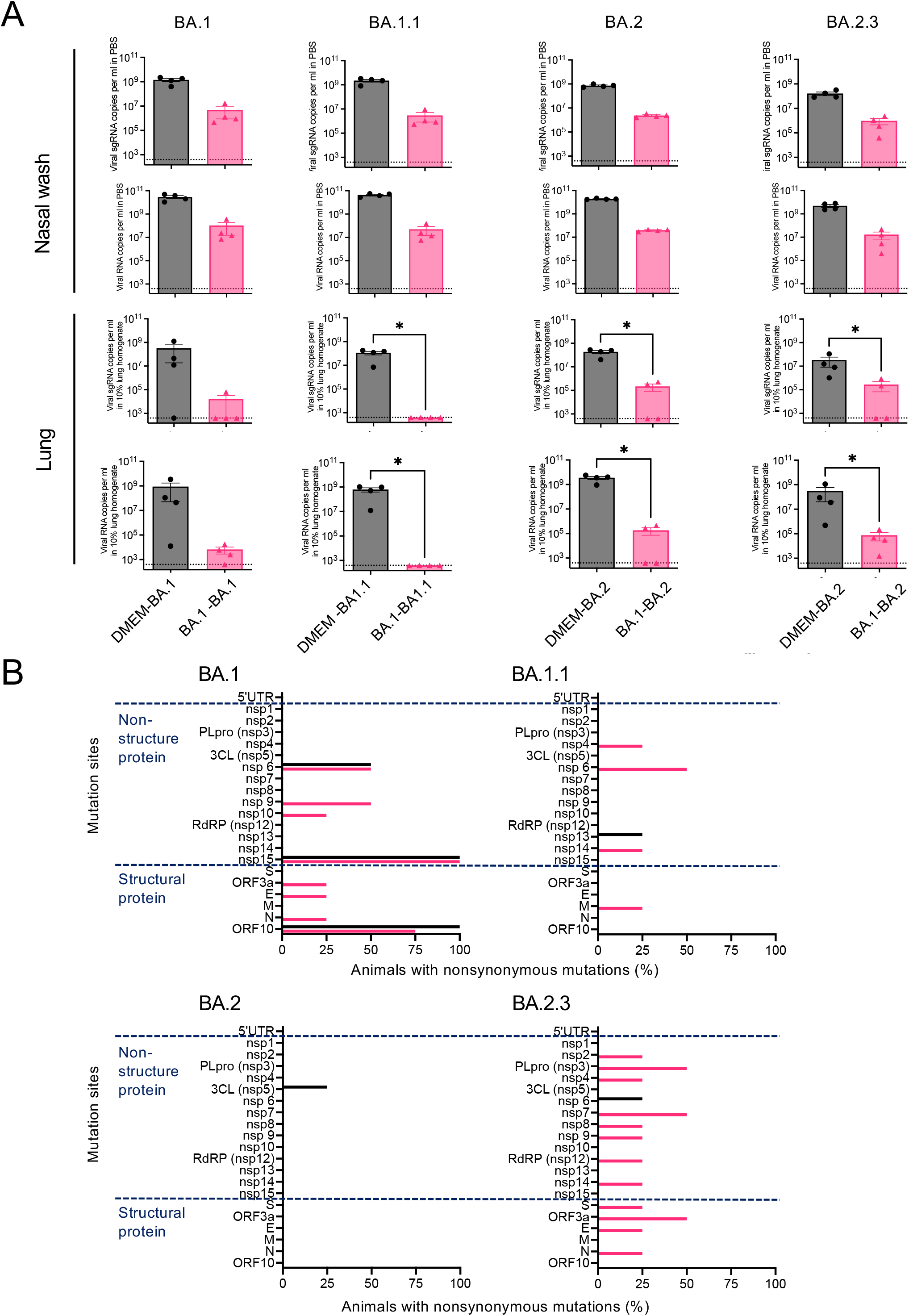
Impact of reinfection in hamsters with Omicron subvariants. Bar graph showing virus-related RNA copies in the nasal wash fluid (upper panels) and supernatant from 10% lung tissue homogenates (lower panels). Dot line indicates detection limit. *, P < 0.05 by Mann-Whitney test. B-D, black bars indicate the primary infection groups and red bars the reinfection groups **(A)**. Percentage of animals with mutations involving amino acid substitutions in each region. Comparison of mutations detected in animals reinfected with SARS-CoV-2. The full-genome sequences of the B. 1.1.529 subvariants recovered from the nasal wash fluid samples (n = 4 per group) were determined using NGS analyses **(B)**.

The full-genome sequence of each of the B.1.1.529 subvariants recovered from the nasal wash fluid samples was determined (Supplementary Table 2 and 3). As in the earlier infection study, nsp15 (ORF1b: K2340T) and ORF10 (ORF10: V30L) nonsynonymous mutations appeared in all animals whose primary infection was with BA.1. (Figure 5B). In reinfected animals, several mutations were detected in nonstructural and structural genes of the variants, though the mutations were unevenly distributed among the infection groups (Figure 5B). Interestingly, for three of the four infection groups fewer within-host variants were detected following primary infection than were detected following reinfection.

In animals from all primary infection groups, pathological lesions consisted of mild to moderate rhinitis and focal broncho-interstitial pneumonia in the respiratory tract associated with virus infection (Figure 6A-D). Animals in the reinfection groups showed very slight to moderate lymphocyte infiltrations in the nasal cavity in the presence or absence of viral antigen (Figure 6A-C). Some animals from the reinfection groups (BA.1-BA.1, one of four; BA.1-BA.1.1, two of four; BA.1-BA.2, two of four in each group) showed focal infiltrations with lymphocytes, plasma cells, and macrophages in the alveoli and around the blood vessels. In reinfected animals, lung pathology occurred even though viral antigen was detected in only one of the animals from the BA.1-BA.2 group (Figure 6A-D). Fibrin deposition was observed in the lungs of most animals after primary infection but in the lungs of very few reinfected animals (Figure 6E). High levels of CXCL10/IP-10 and IL-1β were observed in the nasal wash fluid and lungs of the primary infection groups, but the levels were lower in reinfection animals (Supplementary Figure 2). Thus, exacerbated cytokine production, due to reinfection, was not seen in either the upper or lower respiratory tract. These data suggest that despite the fact that some animals reinfected with subvariants are unable to mount a sufficient immune response against reinfection, especially in the upper respiratory tract, they are still able to eliminate these variants more rapidly, in early phase, than occurs in naïve animals. In addition, there were no findings suggesting disease exacerbation.

**FIG 6.**
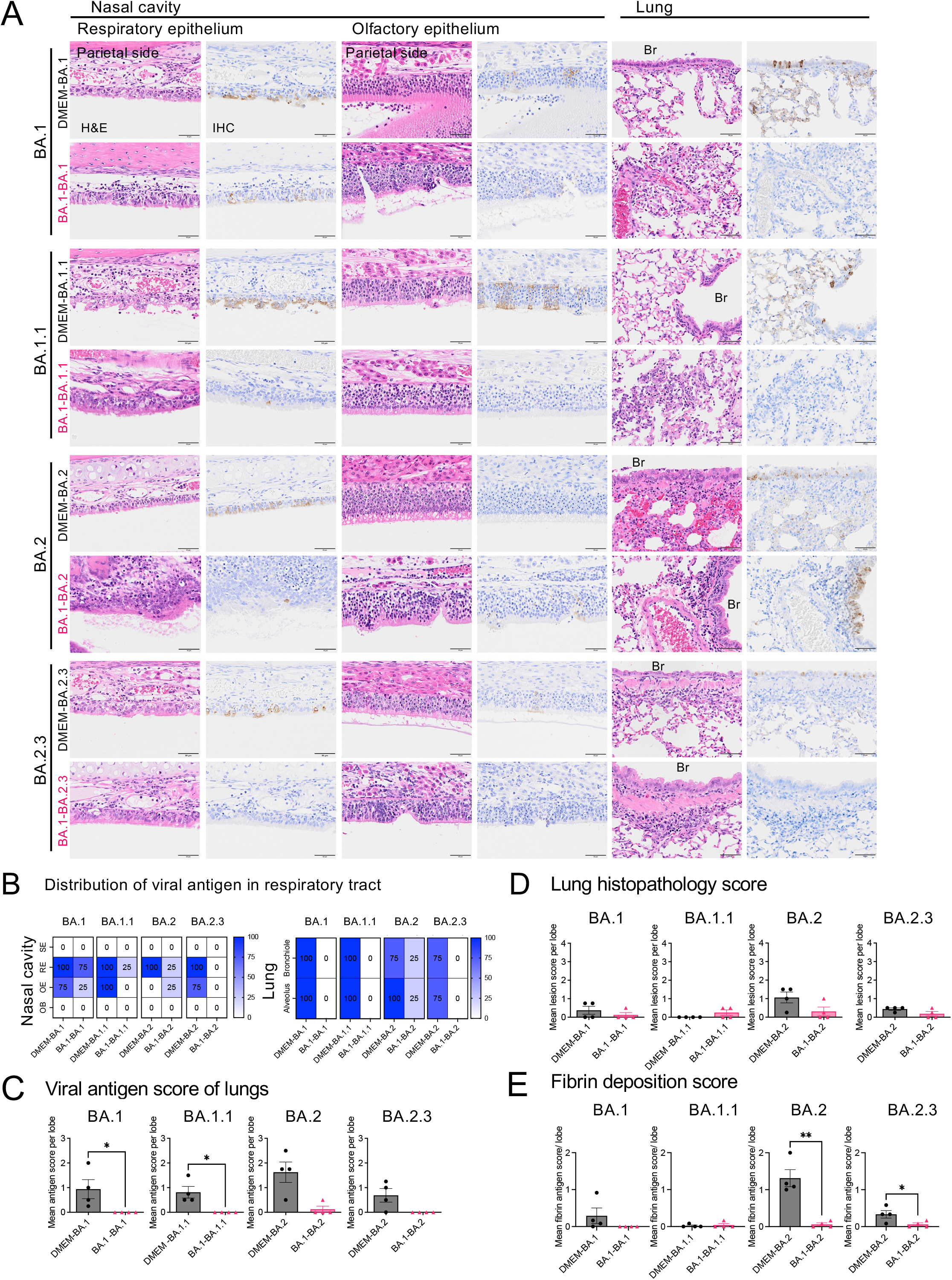
Histopathology of respiratory tract tissues from hamsters following primary- or reinfection with Omicron subvariants. Representative histopathological findings of the nasal cavity and lungs from hamsters after primary or second infection with subvariants of B. 1.1.529 (Omicron) (n = 4) **(A)**. H&E, hematoxylin and eosin staining; IHC, immunohistochemistry for SARS-CoV-2 antigen detection. Br, bronchi. Viral antigen-positive cells were mainly detected in the nasal epithelium on the cranial side. Scale bars, 50 μm. Distribution of viral antigen in respiratory tract from hamsters **(B)**. Heat map shows percentage of viral antigen-positive animals. SE; squamous epithelium, RE; respiratory epithelium, OE; Olfactory epithelium, OB; Olfactory bulb. Viral antigen scores and pathological severity scores of lungs in hamsters **(C and D)**. Fibrin deposition scores of lungs from hamsters by immunohistochemistry. Four lung lobes were taken from each individual animal and scored. *p < 0.05 by one-way ANOVA **(E)**.

## DISCUSSION

Plasma from convalescent human cases and from individuals who had been vaccinated against SARS-CoV-2 exhibited marked reductions in neutralizing activity against Omicron than against the ancestral SARS-CoV-2 (22–26). Multiple lineages of the Omicron variants have emerged including BA.1, BA.2, BA.3, BA.4 and BA.5, with the dominant strain being replaced by an emerging variant (27). Initially, BA.1 was the most prolific sublineage detected worldwide; however, BA.2 is overtaking BA.1 as the dominant variant and now BA. 5 has replaced (27). In human cases, lower neutralizing antibody titers against the BA.4 and BA.5 subvariants than against the BA.1 and BA.2 subvariants suggest that the Omicron variant is continuing to evolve with increasing capacity for neutralization escape (28). Consequently, the monitoring of antigenic changes in new SARS-COV-2 variants that emerge should continue. Syrian hamster models are considered to be ideal for determining the antigenic differences of SARS-CoV-2 variants (29, 30) and, as in a previous study (30), our hamster model showed clear antigenic differentiation between the BA.1 and BA.2 subvariants. Female hamsters were selected for the infection model used in this study since they show better systemic and local antiviral antibody responses and more prolonged humoral immunity, than male hamsters, following SARS-CoV-2 infection (31).

Because the strength of the humoral immune response and the duration of neutralizing antibodies may correlate with disease severity, viral shedding from individuals with mild COVID-19 is a concern (32). In hamsters, primary infection induced immune responses with disease-protective capacity against reinfection by homologous or heterologous Omicron subvariants. In this study, the induction of immunity and neutralizing antibodies, during upper respiratory tract infection, was not a sufficient at preventing proliferation in the upper respiratory tract of hamsters, following reinfection, as reported in previous studies (16, 18, 19). The presence of infectious virus in the upper respiratory tract of hamsters, despite a significant decrease in virus levels following reinfection, suggests that even asymptomatic individuals are capable of shedding infectious virus that can cause infection. On the other hand, it appears that the virus was efficiently cleared from the lower respiratory tract in reinfected animals. Neutralizing antibodies in the blood are believed to be effective in preventing reinfection of the lower respiratory tract (33).

The sequence of variants recovered from hamster nasal wash fluid after reinfection was greater than 99.99% identical to that of the virus used for reinfection. SARS-CoV-2 population in a host is not represented by a single dominant sequence, but rather consists of an ensemble of replicating viruses comprised of closely related sequences called quasispecies (34). Consequently, the alteration of functional genes (such as the S gene) is likely to generate SARS-CoV-2 quasispecies that are better adapted for infection and survival in a particular host (35). Interestingly, viral RNA recovered from nasal washes, from all eight hamsters in two, separate, primary BA.1 infection experiments, showed consistent amino acid substitutions at Nsp15 (ORF1b: K2340T) and ORF10 (ORF10: V30L). By contrast, this consistency was less observed in viral RNA recovered after homologous and heterologous reinfection of Omicron variants, with amino acid substitutions occurring in more diverse regions. Nsp 15 of SARS-CoV-2 functions as an endoribonuclease (36). Nsp15 and several other SARS-CoV-2 proteins inhibit primary interferon production and interferon signaling and thus may interfere with the body’s defense against infection (36–38). Compared with SARS-CoV and other nidoviruses, the Orf10 protein of SARS-CoV-2 is a unique 38 aa protein (39). However, the role of ORF10 is still unclear. Interestingly, the overexpression of ORF10, *in vitro*, markedly suppressed the expression of type I interferon genes (40). In addition, the V30L amino acid substitution in ORF10 correlates with disease severity in COVID-19 patients (41). The significance of these mutations observed in the hamsters remains to be elucidated.

One concern regarding reinfection with the SARS-CoV-2 subtype is that an abnormal immune response to the primary infection may exacerbate the secondary infection and enhance immunopathology (42), as is seen, for example, in antibody-dependent enhancement (ADE) breakthrough infections such as dengue fever (43). Feline infectious peritonitis (FIP), a coronavirus infection in cats, causes classic ADE disease (44, 45); in contrast to FIP, SARS-CoV-2 infections in humans primarily involve the respiratory tract and other organs but not the reticuloendothelial system. It is clear that the primary target of SARS-CoV-2 infection is alveolar epithelial cells, which are directly injured by the virus (46). Since myeloid cells are not the primary target of infection, it is unlikely that vaccine-derived, non-protective, coronavirus antibodies will cause ADE infection in human (47). In addition, no cases of enhanced disease due to reinfection with Omicron have been reported (48–50). CXCL10/IP-10 has a well-established role in the COVID-19-related cytokine storm and is involved in the development of severe lung impairment (51). Elevated levels of key inflammatory chemokines and cytokines, such as CXCL10/IP-10 and IL-1β, were observed in nasal wash fluid and lung homogenate supernatant of hamsters during primary infection, but not after the reinfection. In addition, reinfected hamsters showed no evidence of the eosinophil-associated lung inflammation observed, previously, in infected mice (52). Thus we concluded that pathological and cytokine/chemokine analyses showed no evidence of disease progression after reinfection in our hamster model. However, as new SARS-CoV-2 variants emerge, their virulence could change and, hence, there is a need to actively monitor variants.

Overall, the hamster model demonstrated that immunity acquired during primary infection, with a previous SARS-CoV-2 variant or early Omicron sublineage, suppressed the development of pneumonia after reinfection with a subsequently emerged variant. However, infection and multiplication in the upper respiratory tract after reinfection were inevitable, suggesting the likelihood of virus excretion in asymptomatic hamsters. The virus population in the upper respiratory of reinfected hamsters was shown to be more diverse than that seen in hamsters after primary infection. In addition, differences were observed between the BA.1 strain and other subvariants, in the diversity of virus populations generated in infected hamsters, which may reflect alterations in virus infectivity and replication within specific hamsters. These findings could provide a better understanding of pathology after reinfection by new variants of SARS-CoV-2. In addition, the hamster model should provide insight into the within-host evolution of SARS-CoV-2 and provide important information for countermeasures aimed at diversifying SARS-CoV-2 mutant strains.

## MATERIALS AND METHODS

### Ethics

All procedures involving cells and animals were conducted in a Biosafety Level (BSL) 3 laboratory. All animal experiments were approved by the Animal Care and Use Committee of the National Institute of Infectious Diseases in Japan (approval nos. 120108, 120142, and 121152), and all experimental animals were handled in BSL3 animal facilities according to the guidelines of this committee (approval nos. 19-53, 20-39, and 20-31). All animals were housed in a facility certified by the Japan Health Sciences Foundation.

### Viruses and cells

Viruses were isolated from anonymized clinical specimens (nasopharyngeal/nasal swabs or saliva) collected from individuals diagnosed with COVID-19 as part of the public health diagnostic activities conducted by National Institute of Infectious Diseases (53, 54). VeroE6/TMPRSS2 cells purchased from the Japanese Collection of Research Bioresources Cell Bank (JCRB1819, the National Institute of Biomedical Innovation, Health and Nutrition, Osaka, Japan) were used for viral isolation and viral titrations (53). Cells were cultured in DMEM, low glucose (Sigma-Aldrich, St. Louis, MO), containing 10% FBS, 50 IU/mL penicillin G, and 50 μg/mL streptomycin (10DMEM). Viral infectivity titers were expressed as TCID_50_/mL in VeroE6/TMPRSS2 cells and were calculated according to the Behrens– Kärber method. Work with infectious SARS-CoV-2 was performed under BSL3 conditions.

### Animal experiments

Five-week-old female Syrian golden hamsters (SLC, Shizuoka, Japan) were used for animal experiments. After anesthesia, animals were inoculated intranasally with 1.0×10^3^ TCID_50_ (in 8 μL) of one of five isolates of SARS-CoV-2 including the ancestor strain from lineage A, and four isolates of the VOC (lineage B.1.1.7, B.1.351, B.1.617.2, and B.1.1.529 BA.1; Table 1). All mock-infected hamsters were inoculated with DMEM containing 2% (v/v) FCS containing 50 IU/mL penicillin G, and 50 μg/mL streptomycin (2DMEM). Body weight was measured daily for 3 days (*n* = 4 or 16 per group). Five weeks after their first inoculation, animals were then re-inoculated intranasally with 1.0×10^4^ TCID_50_ (50 μL) of four isolates of the B.1.1.529 subvariants (BA.1, BA.1.1, BA.2, BA.2.3; Table 1). Body weight was measured daily for 3 days (*n* = 3-4 per group), and animals were sacrificed at 3 days p.i. to analyze viral replication and disease pathology (*n* = 3-4 per group). The humane endpoint was defined as the appearance of clinically diagnostic signs of respiratory stress, including respiratory distress and more than 25% weight loss. Animals were euthanized under anesthesia with an overdose of isoflurane if severe disease symptoms or weight loss was observed.

### RNA extraction and quantification of viral RNA genomes

Total RNA from each lung homogenate and nasal wash was isolated using the Maxwell RSC Maxwell RSC Viral Total Nucleic Acid Purification Kit (Promega Corporation), following the manufacturer’s suggested protocol, and quantified by NanoDrop (Thermo Fisher Scientific). The viral RNA copy number in the samples was estimated by real-time RT-PCR. Subgenomic viral RNA transcripts were also detected in N gene transcripts. The primer and probe sets are as follows: NIID_2019-nCOV_N_F2 (5’-AAATTTTGGGGACCAGGAAC-3’), NIID_2019-nCOV_N_R2 (5’-TGGCAGCTGTGTAGGTCAAC-3’), and NIID_2019-nCOV_N_P2 (5’-FAM-ATGTCGCGCATTGGCATGGA-BHQ-3’) for targeting the viral RNA; and SARS2-LeaderF60 (5ʹ-CGATCTCTTGTAGATCTGTTCTCT-3ʹ), SARS2-N28354R (5ʹ-TCTGAGGGTCCACCAAACGT-3ʹ), and SARS2-N28313Fam (5ʹ-FAM-TCAGCGAAATGCACCCCGCA-TAMRA-3ʹ) for targeting the subgenomic RNA. The reaction mixtures were incubated at 50°C for 30 min, followed by incubation at 95°C for 15 min, and thermal cycling, which consisted of 40 cycles of denaturation at 94°C for 15 s, and annealing and extension at 60°C for 60 s. This assay was performed on a LightCycler 480 (Roche, Basel, Switzerland).

### SARS-CoV-2 neutralizing assay

Blood was obtained from each hamster under anesthesia (55) and when euthanized. Sera were then obtained by centrifugation and were inactivated by incubation at 56°C for 30 min. Aliquots (100 TCID_50_/well) of SARS-CoV-2 were incubated at 37°C for 1 h in the presence or absence of hamster serum (serially diluted two-fold), and then added to confluent VeroE6/TMPRSS2 cell cultures in 96-well microtiter plates. Samples were examined for viral cytopathic effects on Day 5, and the neutralizing antibody titers were determined as the reciprocal of the highest dilution at which no CPEs were observed. The lowest and highest serum dilutions tested were 1:16 and 1:2048, respectively.

### ACE2 binding inhibition electrochemiluminescence immunoassay

A multiple assay for neutralizing antibodies to spike antigens from variants of SARS-CoV-2 using the V-PLEX SARS-CoV-2 (ACE2) kits (K15570U and K15586U, Meso Scale Discovery) was used. 1:10 and 1:100 diluted sera were used for the assay. The assay samples were read on a high-performance electrochemiluminescence immunoassay instrument, MESO QuickPlex SQ 120 (Meso Scale Discovery), as described by the manufacturer.

### Detection of inflammatory cytokines and chemokines

Homogenized lung tissue samples (10% w/v) and nasal wash samples were diluted 1:1 in cell extraction buffer (10 mM Tris, pH 7.4, 100 mM NaCl, 1 mM EDTA, 1 mM EGTA, 1 mM NaF, 20 mM Na_4_P_2_O_7_, 2 mM Na_3_VO_4_, 1% Triton X-100, 10% glycerol, 0.1% SDS, and 0.5% deoxycholate (BioSource International, Camarillo, CA)), incubated for 10 min on ice with vortexing, irradiated for 10 min with UV-C light to inactivate infectious virus, and tested in the BSL2 laboratory. Cytokine and chemokine levels were measured with a commercial rat cytokine/chemokine magnetic bead panel 96-well plate assay kit (Milliplex MAP kit, Merck Millipore), which detects 5 cytokines and chemokines including IP-10/CXCL10, IL-1α, IL- 1β, IL-10, and VEGF (56). The assay samples were read on a Luminex 200 instrument with xPONENT software (Merck Millipore), as described by the manufacturer.

### Histopathology and immunohistochemistry

The lungs and head including nasal cavity and brain were harvested and fixed in 10% phosphate-buffered formalin. Fixed tissues were routinely embedded in paraffin, sectioned, and stained with hematoxylin and eosin (H&E). For immunohistochemistry, antigen retrieval of the formalin-fixed tissue sections was performed by autoclaving at 121°C for 10 min in retrieval solution at pH 6.0 (Nichirei, Tokyo, Japan). SARS-CoV-2 antigens were detected using a polymer-based detection system (Nichirei-Histofine Simple stain MAX PO; Nichirei Biosciences, Inc., Tokyo, Japan), and an in-house rabbit anti-SARS-CoV-2 N antibody was used as the primary antibody. Nuclei were counterstained with hematoxylin for 10 s.

Histopathology scores were determined based on the percentage of lesion area including inflammation, hemorrhage and edema, as determined by HE staining in each group by using the following scoring system: 0, no lesion; 1, focal lesion within 30% or less total area; 2, diffuse lesion involving 30–70% total area; 3, diffuse lesion involving more than 70% total area. Scores were also determined based on the percentage of virus antigen-positive cells, as determined by immunohistochemistry in each group by using the following scoring system: 0, no antigen-positive cells; 1, antigen-positive cells were occasionally observed in each cut sections (1–3 antigen-positive areas per section were observed in the high magnification); 2, scattered positive cells were observed (4–9 antigen-positive areas per section were observed in the high magnification); 3, many positive cells were diffusely and widely observed (more than 10 antigen-positive areas per section were observed in the low magnification). Mean scores from all lung sections (four lung sections/animal) in each animal were calculated. Dots in figure indicate mean scores in each animal.

### Next generation sequencing analysis for comparison of SARS-CoV-2 mutations

To identify major population of virus in respiratory tract of hamsters after reinfection, a next generation sequencer was used to obtain the entire length of the viral genome. The sequences obtained from the samples were compared with those of the inoculated viruses. The viral RNAs were extracted from the homogenized lung tissue samples (10% w/v) and nasal wash samples using the Maxwell® RSC Viral Total Nucleic Acid Purification Kit (Promega, Madison, WI). The whole genomes of SARS-CoV-2 used in this research were amplified using a modified ARTIC protocol with several primers replaced or added (57, 58). The viral cDNAs were synthesized from extracted RNA using the Luna Script RT Super Mix Kit (New England BioLabs, Ipswich, MA), followed by DNA amplification by multiplex PCR in two separated primer pools using ARTIC-N5 primers (59, 60) and Q5 Hot Start DNA polymerase (New England BioLabs). The DNA libraries for Illumina NGS were prepared from pooled amplicons using the QIAseq FX DNA Library Kit (QIAGEN) and analyzed using the iSeq 100 and MiSeq (Illumina). The obtained reads were analyzed by the CLC Genomics Workbench (version 21, QIAGEN) with the Wuhan/Hu-1/2019 sequence (GenBank accession number MN908947) as a reference. The sequence data have been deposited in the DNA Data Bank of Japan (DDBJ) Sequence Read Archive, under submission (BioProject Accession: PRJDB14262; BioSample accessions: SAMD00523210-SAMD0052326). In the reinfection experiment, the frequencies of gene mutations were calculated with each protein. The average depth of aligned reads was 3132.

In the co-infection experiment, the ratio of Delta and BA.1 was calculated as the percentages based on the 10 regions where these two viruses can be distinguished. The nucleotide numbers of these 10 regions are: 2832, 8393, 11537, 21618, 22673-22674, 23048, 23063, 23604, 26530, and 28311 of the Wuhan/Hu-1/2019 genome, which correspond to amino acids 856, 2710, 3758 in ORF1a, 19, 371, 496, 501, 681 in the spike protein, 3 in the M protein, and 13 in the N protein, respectively. The ratio of BA.1 and BA.2 was also analyzed based on the 10 regions where these two strains could be identified. The nucleotide numbers in these 10 regions are: 2832, 8393, 11537, 21618, 22204-22205, 22673-22674, 22786, 22898, 23048, and 29510 in the Wuhan/Hu-1/2019 genome, which correspond to amino acids 856, 2710, 3758 in ORF1a, 19, 214, 371, 408, 446, 496 in the spike protein, and 13 in the N protein, respectively. The average depth of aligned reads was 3132. Each read depth at the 20 regions used to calculate the ratios was more than 200.

### Statistical analysis

All data are expressed as the mean and standard error of the mean, except for neutralizing antibodies (Geometric mean titers with 95% confidence interval, GMT+95%Cl). Statistical analyses were performed using GraphPad Prism 9 software (GraphPad Software, La Jolla, CA). Intergroup comparisons were performed using nonparametric analysis. A P value < 0.05 was considered statistically significant.

## ACKNOWLEDGMENTS

We thank Dr Shutoku Matsuyama and Dr Makoto Takeda (National Institute of Infectious Disease) for providing SARS-CoV-2 isolates. We are grateful to Midori Ozaki, Takiko Yoshida, and Dai Izawa for their technical assistance and our colleagues at the Institute for helpful discussions. We thank the members of the Management Department of Biosafety and Laboratory Animals for support with the BSL3 facility.

This work was supported by the Grant-in-Aid for Scientific Research from the Ministry of Education, Culture, Sports, Science, and Technology in Japan (21K20767 to NS-S; 20K21666 to NN) and the Japan Agency for Medical Research and Development grants (JP21fk0108615 to KM and NN; JP21nf0101626 to HH; JP21wm0125008 to TS).

---

**Supplementary Figure 1.**
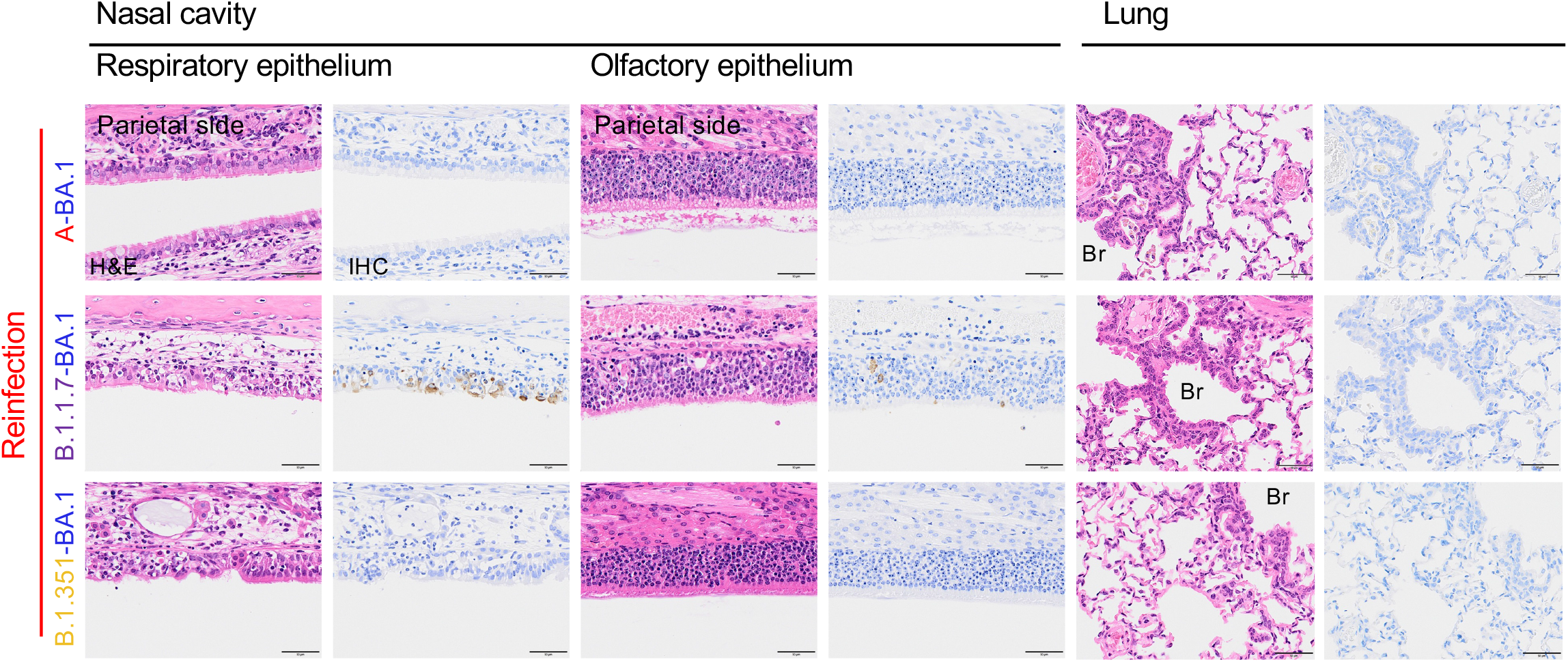
Histopathology of respiratory tract tissues after reinfection with an Omicron BA.1 variant. Representative histopathological findings of the nasal cavity and lungs from hamsters 3 days after second infection with an isolate of TY38-873, B. 1.1.529 (Omicron) subvariant BA.1 (n = 3 or 4). H&E, hematoxylin and eosin staining; IHC, immunohistochemistry for SARS-CoV-2 antigen detection. Br, bronchi. Viral antigen-positive cells were mainly detected in the nasal epithelium on the cranial side. Scale bars, 50 μm.

**Supplementary Figure 2.**
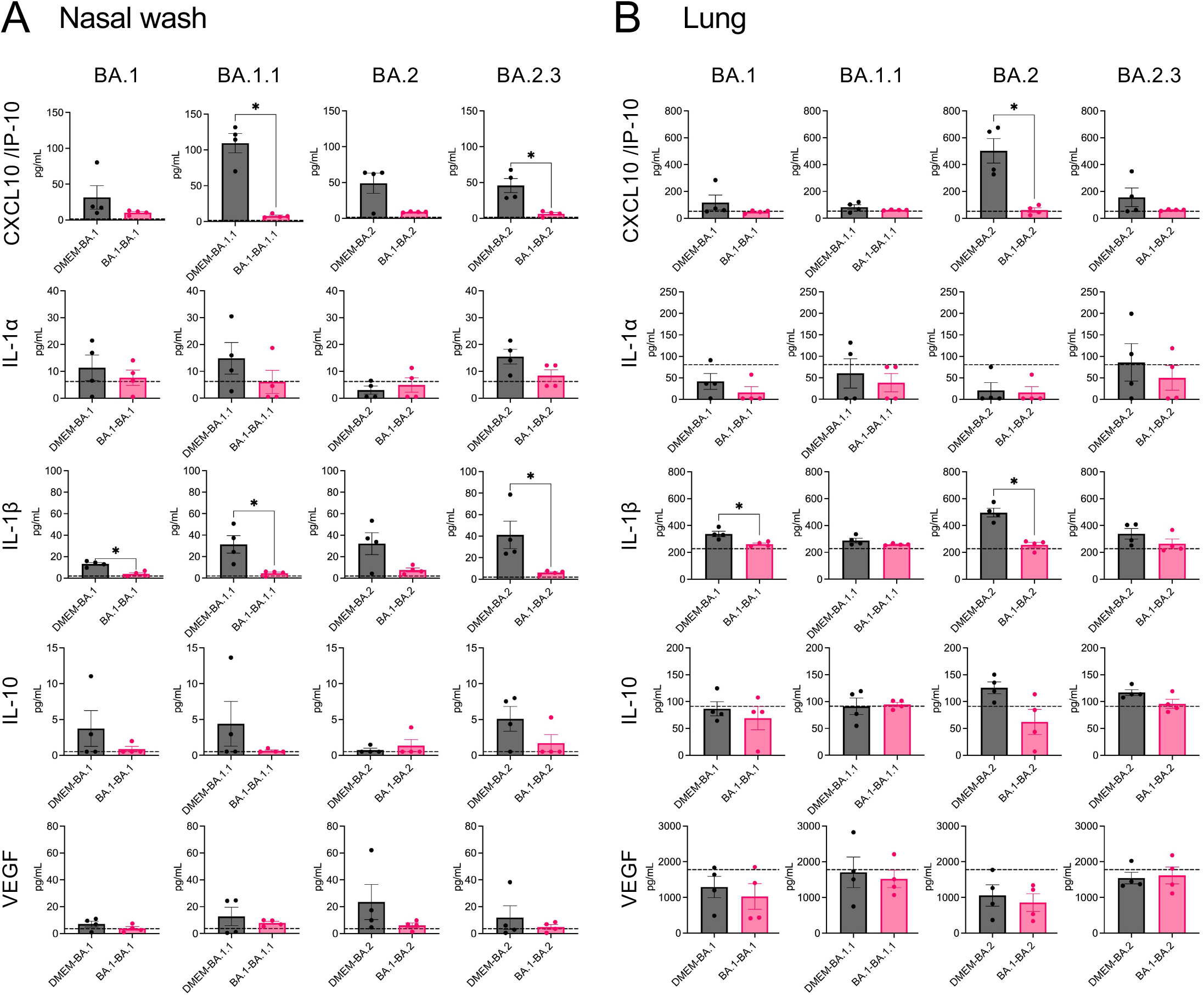
Cytokine and chemokine levels in the respiratory airway of hamsters 3 days post primary or reinfection. The nasal wash fluid **(A)** and supernatant from 10% lung tissue homogenates **(B)** of hamsters at 3 days p.i. (n = 4) *, P < 0.05 by Mann-Whitney test. Black bars indicate the first infection group; red bars, reinfection group. Dashed line indicates mean values in the nasal wash fluid or lung homogenates from mock-infected hamsters (n = 4, at 3 days p.i. with DMEM).

**Table S1.**
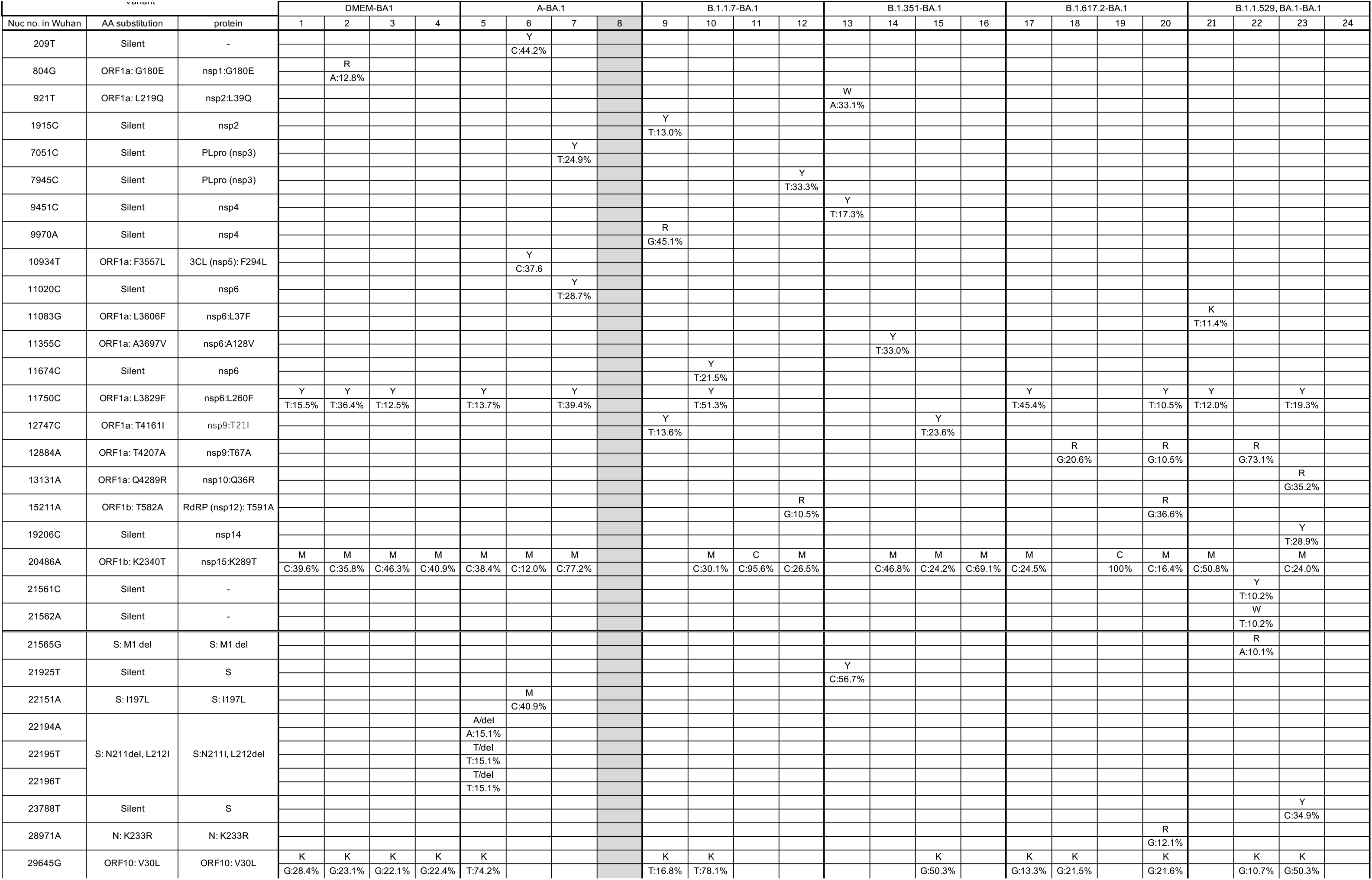

**Table S2.**
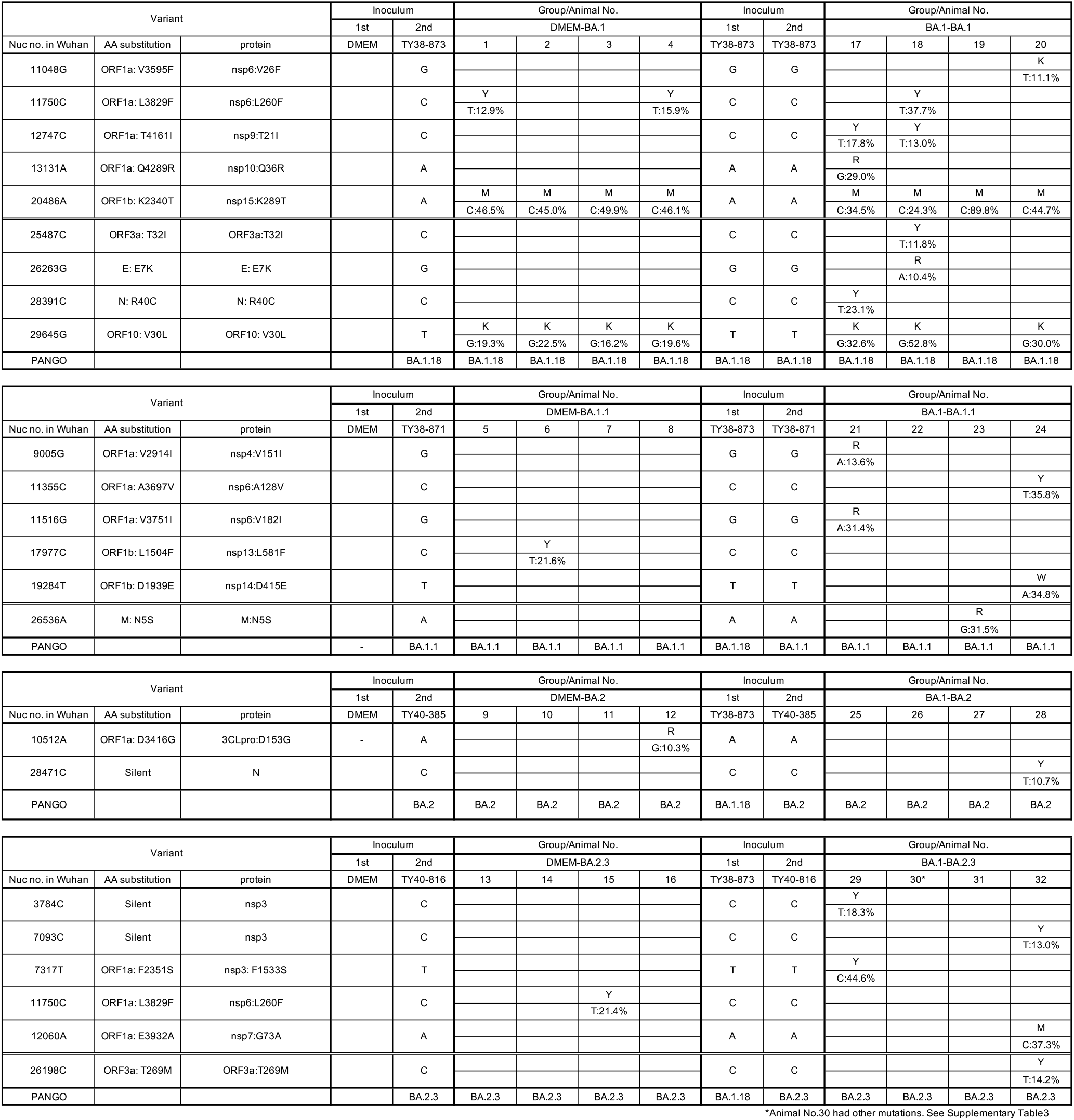

**Table S3.**

|  |  |  | Inoculum |  | Group/Animal No. |
| --- | --- | --- | --- | --- | --- |
| 14-3, ham#29-32<br>1st: TY38-873, 2nd: TY40-816 |  |  | 1st | 2nd | BA.1-BA.2.3 |
| Nuc no. in Wuhan | AA substitution | Protein | TY38-873 | TY40-816 | 30 |
| 2086G | ORF1a: Q607H | nsp2: Q427H | G | G | K<br>T:10.9% |
| 4250T | ORF1a: Y1329H | nsp3: Y511H | T | T | Y<br>C:15.2% |
| 5514T | ORF1a: V1750A | nsp3: V932A | T | T | Y<br>C:15.9% |
| 5763G | ORF1a: C1833F | nsp3: C1015F | G | G | K<br>T:10.9% |
| 8600G | ORF1a: V2779L | nsp4: V16L | G | G | S<br>C:12.8% |
| 12054T | ORF1a: C3931fs | nsp7: C72fs | T | T | del<br>10.3% |
| 12467G | ORF1a: A4068T | nsp8: A126T | G | G | R<br>A:10.8% |
| 12708C | ORF1a: A4148V | nsp9: A8V | C | C | Y<br>T:10.1% |
| 14460T | ORF1b: F331L | nsp12: F340L | T | T | K<br>G:22.3% |
| 18352T | ORF1b: L1629del,<br>P1630del | nsp14: L105del,<br>P106del | T | T | del<br>13.4% |
| 18353T |  |  | T | T | del<br>13.4% |
| 18354A |  |  | A | A | del<br>13.4% |
| 18355C |  |  | C | C | del<br>13.4% |
| 18356C |  |  | C | C | del<br>13.4% |
| 18357T |  |  | T | T | del<br>13.4% |
| 18786T | Silent | nsp14 | T | T | Y<br>C:10.3% |
| 21575C | S: L5F | S: L5F | C | C | Y<br>T:11.4% |
| 23766C | S: S735L | S: S735L | C | C | Y<br>T:11.4% |
| 25613C | ORF3a: S74F | ORF3a: S74F | C | C | Y<br>T:13.0% |
| 25714C | ORF3a: L108F | ORF3a: L108F | C | C | Y<br>T:12.3% |
| 26355T | Silent | E | T | T | Y<br>C:13.2% |
| 26413T | E: Y57H | E: Y57H | T | T | Y<br>C:12.8% |
| 28864A | Silent | N | A | A | W<br>T:18.9% |
| 29409C | N: T379N | N: T379N | C | C | M<br>A:16.1% |
| 29708C | Silent | 3'UTR | C | C | Y<br>T:11.2% |
| PANGO |  |  | BA.1.18 | BA.2.3 | BA.2.3 |

